# Altered visual population receptive fields in human albinism

**DOI:** 10.1101/726141

**Authors:** Ivan Alvarez, Rebecca Smittenaar, Sian E. Handley, Alki Liasis, Martin I. Sereno, D. Samuel Schwarzkopf, Chris A. Clark

## Abstract

1

Albinism is a congenital disorder where misrouting of the optic nerves at the chiasm gives rise to abnormal visual field representations in occipital cortex. In typical human development, the left occipital cortex receives retinal input predominantly from the right visual field, and vice-versa. In albinism, there is a more complete decussation of optic nerve fibers at the chiasm, resulting in partial representation of the temporal hemiretina (ipsilateral visual field) in the contralateral hemisphere. In this study, we characterize the receptive field properties for these abnormal representations by conducting detailed fMRI population receptive field mapping in a rare subset of participants with albinism and no ocular nystagmus. We find a nasal bias for receptive field positions in the abnormal temporal hemiretina representation. In addition, by modelling responses to bilateral visual field stimulation in the overlap zone, we found evidence in favor of discrete unilateral receptive fields, suggesting a conservative pattern of spatial selectivity in the presence of abnormal retinal input.

**Highlights:**

- We characterized population receptive fields in albinotic participants with no ocular nystagmus
- Confirmed overlapping representations of left and right visual fields in cortex
- Detected nasal bias in receptive field location for temporal hemiretina representation
- Evidence in favor of discrete unilateral receptive fields and against double coding

## 2 Introduction

Albinism is a congenital disorder associated with misrouting of the optic nerves during embryogenesis, which leads to abnormal retinotopic organization in sub-cortical and cortical visual areas (Carroll, Jay, McDonald, & Halliday, 1980; Creel, Witkop, & King, 1974; Hedera et al., 1994; Morland, Hoffmann, Neveu, & Holder, 2002). In humans, retinal projections are normally divided at the optic chiasm, with temporal hemiretina fibers projecting to the hemisphere ipsilateral to the eye, and nasal hemiretina fibers crossing the midline and projecting to the contralateral hemisphere. In albinism, however, the line of decussation is shifted, leading to an over-crossing of temporal hemiretina projections to the contralateral hemisphere and leaving a weaker ipsilateral projection (Guillery, Okoro, & Witkop, 1975; Neveu & Jeffery, 2007). Despite this gross anatomical abnormality, individuals with albinism have relatively normal visual spatial perception and behavior, aside from peripheral effects such as reduced acuity as a result of foveal hypoplasia (Kinnear, Jay, & Witkop, 1985; Summers, 1996).

The functional consequences of this misrouting are not fully understood. It is clear from studies in the cat that a distinct temporal visual field representation forms in the contralateral lateral geniculate body (Hubel & Wiesel, 1971) and visual cortex (Kaas & Guillery, 1973) and the topology of the abnormal representation is variable (Cooper & Blasdel, 1980). Three models of functional organization in mammalian albinism have been proposed; (1) a contiguous representation, where ipsilateral and contralateral visual fields form a continuous map, (2) an interleaved representation, where mirror-symmetric ipsilateral and contralateral visual field locations are represented on the same cortical territory and (3) an interleaved suppressed representation, where input to the abnormally-routed representation is suppressed (Guillery, 1986; Guillery, Casagrande, & Oberdorfer, 1974; Hoffmann & Dumoulin, 2015; Kaas, 2005; Shatz & LeVay, 1979). However, these models are largely derived from experiments in the cat, and evidence for their applicability in primates is limited. Nevertheless, a single case study in a green monkey (Guillery *et al.*, 1984) and more recently human functional MRI (fMRI) studies (Hoffmann, Tolhurst, Moore, & Morland, 2003; Kaule et al., 2014; Morland et al., 2002) suggest the presence of overlapping representations of ipsilateral and partial contralateral visual fields on the same cortical territory, in agreement with model (2); an interleaved representation. While these recent advances shed light on the topographic organization of the albinotic visual cortex, relatively little is known about the nature of these abnormal representations.

First, it is unclear if the abnormal contralateral representation has a coarse or fine sampling of the visual field. If coarse, it might merely increase spatial integration. If fine, it might support a duplicate coding of fine spatial detail within the same hemisphere (Kaas, 2005). Second, the neural encoding model for overlapping receptive fields remains poorly understood. One interpretation of the interleaved pattern is that two maps representing different parts of the visual field overlap, resulting in ipsilateral and contralateral representations within the same cortical territory, perhaps organized in hemifield columns (Guillery, 1986; Guillery *et al.*, 1984). However, the normal integration across adjacent columns would be confounded by the dual, offset visual field representations, and would not obviously support inter-ocular integration for standard stereoscopic vision (Klemen, Hoffmann, & Chambers, 2012). Instead, integrating neurons might have ipsilateral and contralateral receptive fields - a “dual receptive field”. Alternatively, binocular cells in early visual cortex may be solely modulated by classical receptive field cells capturing sensory input from a single hemifield. Such integration cells would require substantial intracortical plasticity in order to suppress hemifield crosstalk and give rise to integrated stereoscopic vision. The presence or absence of evidence for dual receptive fields would therefore inform how abnormal retinal input is integrated to produce stereoscopic vision and to what degree are retinotopic representations plastic in the abnormally developed visual cortex.

In order to characterize receptive field properties in human albinism, we used a fMRI population receptive field (pRF) mapping approach in a rare sub-population of participants with albinism and no nystagmus. Consistent fixation is critical for delivering visual stimulation in a retinotopic fashion, as the position of the stimulus on the retina must be known in order to correctly reconstruct activity patterns in cortex (Dumoulin & Wandell, 2008; Wandell & Winawer, 2015). In the presence of involuntary eye movements, the correspondence between the stimulus presented and the retinal image cannot be ensured. Thus, fixation stability is a concern for visuotopic mapping in general (Binda, Thomas, Boynton, & Fine, 2013) and pRF mapping in particular (Hummer et al., 2016; Levin, Dumoulin, Winawer, Dougherty, & Wandell, 2010). As nystagmus is a primary clinical feature of albinism, present in 89-95% of albinism cases (Apkarian & Shallo-Hoffmann, 1991; Charles, Green, Grant, Yates, & Moore, 1993; Lee, King, & Summers, 2001), it presents a significant challenge for delivering retinotopic stimulation in the presence of involuntary eye movements. In light of these considerations, we have opted to study a small group of individuals with clinically albinotic phenotypes, but not presenting with overt nystagmus and consistent fixation behavior.

Following monocular stimulation of the temporal hemiretina, significant blood-oxygen level dependent (BOLD) responses were observed in contralateral occipital cortex, demonstrating retinotopic organization within the temporal hemiretina representation largely overlapping the nasal representation, in agreement with the interleaved representation model. We report no differences in pRF size for nasal or temporal hemiretina representation in the shared territory of V1, V2 and V3, pointing to similar computational roles for the overlapping representations. We detected a nasal bias for pRF position in the abnormal temporal hemiretina representation. Finally, by modelling responses to bilateral visual field stimulation we found no evidence in favor of a dual receptive field model. Instead, our data points to interleaved and unsuppressed representations of both nasal and temporal hemiretina, occupying the same cortical territory (model 2), suggesting a conservative pattern of functional organization in early visual areas under grossly abnormal retinal input.

## 3 Materials and methods

### 3.1 Participants

We report how we determined our sample size, all data exclusions (if any), all inclusion/exclusion criteria, whether inclusion/exclusion criteria were established prior to data analysis, all manipulations, and all measures in the study. Five participants with albinism (A1-A5) took part in the study (2 females, mean age=23.80, *SD*=15.90, age range=9-50). A diagnosis of albinism was reached after an ophthalmological assessment that investigated the presence or absence of phenotypic markers. These markers included the following: iris transillumination, photophobia, skin color, hair color at birth, hair color on the day of assessment, and visual appearance of macula. Retinal imaging was performed in a clinical setting, revealing foveal hypoplasia. Finally, the presence or absence of a crossed asymmetry was assessed by comparison of monocular visual evoked potential (VEP) distribution in a clinical test.

Based on their phenotypic features, A1, A4 and A5 were diagnosed with oculocutaneous albinism (OCA), and A2 and A3 were diagnosed with ocular albinism type OA1. A4 and A5 are full siblings. An overview of the key phenotypic description is provided in 0. No genotype information was available at the time of testing.

In addition to participants with albinism, 10 healthy adult controls (4 males, mean age=26.30, SD=4.95, age range: 21-36) with normal visual acuity took part in the study. No exclusions were made, with all participants recruited (albinotic and control) included in all analysis and results presented.

All adult participants provided written informed consent. Parents or legal guardians provided written consent on behalf of underage participants. This study was approved by the London - City and East Research Ethics Committee of the UK Health Research Authority.

**Table 1.** Phenotypic markers for participants with albinism. A-Type = Type of albinism, BMVA = best monocular visual acuity (LogMAR), ITI = Iris transillumination, MA = Macula appearance, VEP-X-Assy = VEP crossed asymmetry. A1, A4 and A5 were diagnosed with oculocutaneous albinism (OCA) after presenting with bilateral iris transillumination and blonde fundus. A2 and A3 were diagnosed with ocular albinism (OA1) and presented regular fundus pigmentation and minimal iris transillumination. Participants with albinism displayed VEP asymmetry and foveal hypoplasia consistent with a shift of the line of decussation.

|  | Sex | Age | A-Type | BMVA | Stereopsis | ITI | MA | Fundus | VEP-X-assy |
| --- | --- | --- | --- | --- | --- | --- | --- | --- | --- |
| **A1** | F | 21 | OCA | 0.140 | no | bilateral | granular | blonde | yes |
| **A2** | M | 25 | OA1 | 0.240 | no | minimal | granular | regular pigmentation | yes |
| **A3** | F | 50 | OA1 | 0.240 | no | minimal | granular | regular pigmentation | yes |
| **A4** | M | 9 | OCA | 0.260 | no | bilateral | granular | blonde | not tested |
| **A5** | M | 14 | OCA | 0.200 | no | bilateral | granular | blonde | yes |

### 3.2 Ophthalmological screening

All participants underwent ophthalmological screening prior to taking part in the study. Best corrected monocular visual acuity was assessed with an ETDRS chart at 4 m distance to threshold. The best monocular acuity for the control group was M=0.03, SD=0.21 LogMAR, and all participants with albinism performed at a minimum of 0.3 LogMAR. All fMRI stimulation was delivered monocularly to the best-performing eye. All five participants with albinism failed to demonstrate binocular stereopsis when assessed with the Frisby stereotest (Frisby Near Stereotest, Sheffield, UK).

Visual field perimetry was assessed with automated perimetry equipment (Octopus 900, Haag-Streit Diagnostics, Koeniz, Switzerland) to two standard isopters (outer target size=I4e, inner target size=I2e, target speed=5°/s). Resulting visual fields showed normal outer perimeter detection for A1-A5 (see Appendix A). Inner perimeter detection showed a small reduction in the nasal visual field of A2 and the temporal visual field of A4.

Optical imaging of the retina was performed with a spectral domain OCT (optical coherence tomography) device (SPECTRALIS, Heidelberg Engineering, Heidelberg, Germany) with active eye tracking (axial resolution=7μm, lateral resolution=14μm, scan rate=40 kHz, scan depth=1.8 mm, field of view=9 mm, averages=36). A1-A5 showed an absent foveal pit and bilateral foveal hypoplasia, characteristic of albinism (see Appendix B).

Consistent eye fixation is a requirement for visuotopic mapping, particularly if fixation behavior differs between two groups being investigated (Bressler & Silver, 2010; Crossland, Morland, Feely, Hagen, & Rubin, 2008). We assessed the stability of eye movement at fixation outside the scanner for participants with and without albinism to establish whether differences were expected during fMRI acquisition. Fixation was assessed monocularly, in the eye with best monocular acuity, that is, the eye to which monocular stimulation was delivered during the fMRI experiment.

Eye movement recordings were carried out with a head-mounted infrared eye tracker (JAZZ-novo, Ober Consulting, Poznan, Poland) sampling relative eye position during visual stimulation on a plasma display. First, a 1.15° fixation target was presented for 10 s in two locations, 14° to left or to the right from central fixation for calibration. A total of 5 events were presented in each position. Next, a central fixation target was presented for 300 s, with the participant instructed to maintain a constant head position and fixate to the targets as they appeared. In order to quantify stability at central fixation, horizontal axis traces were visually inspected, and blink artefacts identified and removed at their amplitude peak. Time points 200 ms immediately before and after blink peaks were also removed from the analysis. Resulting traces were then de-trended to remove linear drifts introduced by the apparatus. Standard deviations of horizontal axis displacement for the eye with best monocular acuity during central fixation were 0.57° for A1, 0.86° for A2, 0.84° for A3, 0.59° for A4, and 0.81° for A5. Fixation stability in the control group averaged 0.64° (95% CI=0.15° - 1.13°). Participants with albinism showed comparable horizontal stability to the control group, with no systematic deviation of gaze at central fixation.

### 3.3 fMRI visual stimulation

Stimuli were generated in MATLAB (v8.0, Mathworks Inc., Natick, MA, USA) using Psychtoolbox (Brainard, 1997; Pelli, 1997) and publicly available at https://osf.io/3mf6n. Stimuli were displayed on a back-projection screen in the bore of the magnet via an LCD projector. This arrangement ensured the peripheral visual field was stimulated.

The stimulus pattern consisted of a 62° radius disc of a dynamic, high-contrast tessellated pseudo-checkerboard with a drifting ripple-like pattern that varied across time in spatial frequency (Alvarez, de Haas, Clark, Rees, & Schwarzkopf, 2015). This pattern was presented in two configurations; a hemifield mapping stimulus for pRF mapping and full-field configuration for estimation of the subject-specific hemodynamic response function (HRF). Stimuli were delivered monocularly to the eye with best-uncorrected visual acuity, while the poorer-performing eye was covered with an eyepatch. The hemifield mapping stimulus consisted of the pattern described, displayed on a single hemifield as divided by the vertical meridian, either ipsilateral or contralateral to the occluded eye, and presented through two simultaneous ‘wedge’ and ‘ring’ apertures on an equiluminant grey background (see Figure 1). A wedge section of 18° of the stimulus circumference rotated either clockwise or counter-clockwise along the polar dimension while a truncated ring section expanded or contracted, scaling between 0.66° and 27.54° eccentricity. Crucially, neither aperture extended beyond the vertical meridian. The maximum stimulated position for all participants was 62° from fixation.

**Figure 1.**
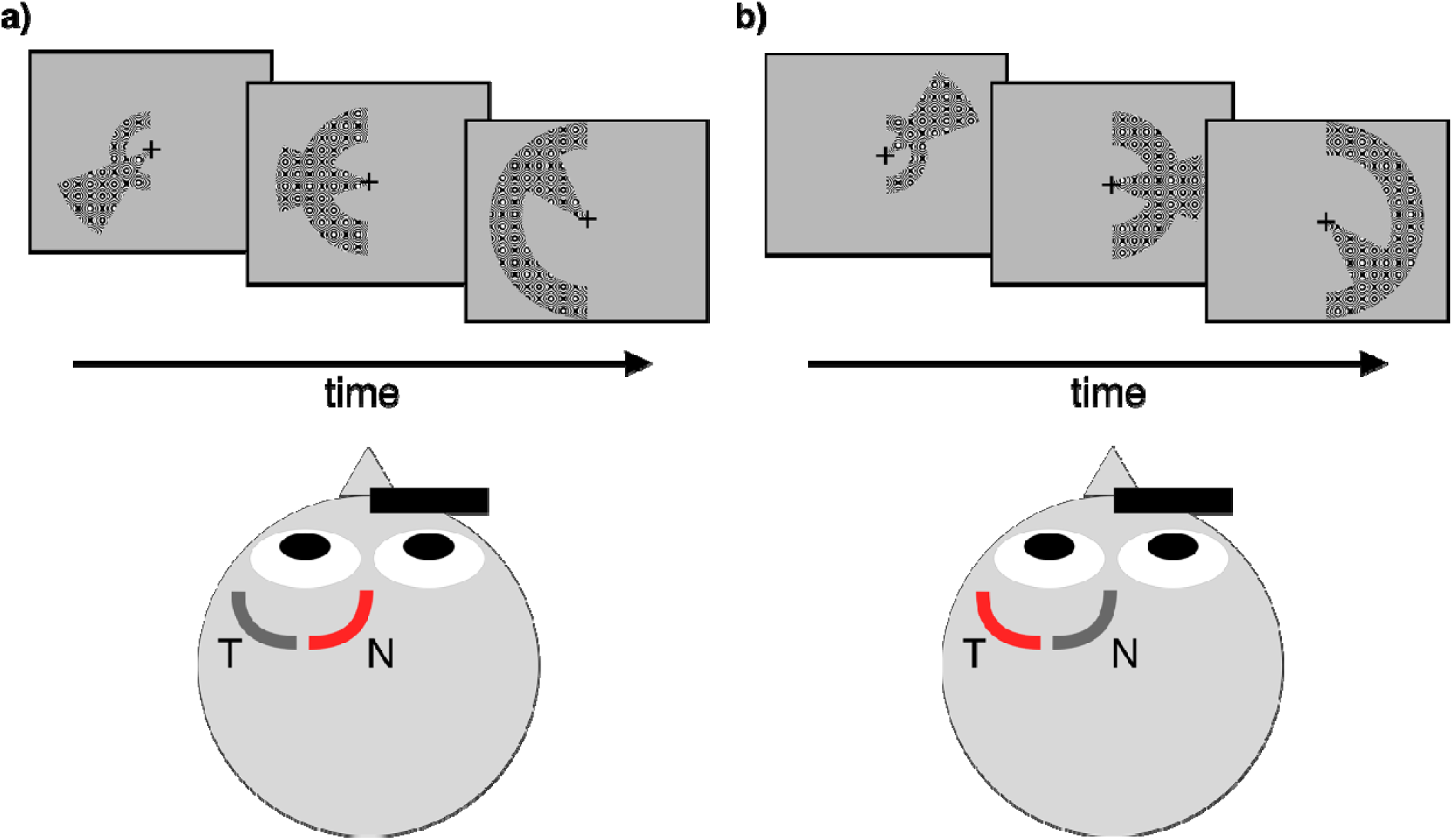
Experimental design. A retinotopic mapping stimulus consisting of simultaneous and de-phased, contrast-reversing wedge and ring apertures was presented monocularly to either (a) left or (b) right hemifield, divided by the vertical meridian. The eye with best uncorrected monocular acuity was stimulated, with left depicted here for illustration. Two runs were acquired for each hemifield, cycling in opposite directions and each followed by a mean luminance period.

Apertures presented cycled at different frequencies, with the wedge and ring apertures completing a cycle every 56 s and 40 s, respectively. Apertures changed position every 1 s on the onset of each EPI volume acquired. A single run of the stimulus consisted of 5 wedge and 7 ring revolutions, followed by 30 volumes of equiluminant grey background, totaling 310 volumes acquired per run. A total of 4 runs were conducted, two presented ipsilateral and two contralateral to the occluded eye. Each pair contained one run with clockwise wedge rotation and expanding rings, and one run with counter-clockwise wedge rotation and contracting rings. The order of presentation was randomized across participants, with a total of 1240 volumes acquired per participant.

The full field configuration consisted of the stimulus pattern presented through a 62° radius circular aperture for 3 volumes, followed by 32 volumes of equiluminant grey background. This was repeated 10 times, totaling 350 volumes acquired.

Throughout both conditions, a fixation cross spanning 1.8° was presented to aid fixation. Participants engaged in an attentional task, consisting of a brief (200 ms) change in color of the fixation cross, occurring semi-randomly with a probability of 5% for any given 200 ms epoch, with no consecutive events. Participants were instructed to attend to the cross and provide a response via an MRI-compatible response button every time they witnessed an event. Participant responses were monitored to ensure engagement with the task.

### 3.4 MRI acquisition and pre-processing

MR images were acquired on a 1.5T Avanto MRI system using a 32-channel head coil (Siemens Healthcare, Erlangen, Germany). During visual stimulation, the top elements of the head coil were removed (remaining coils=20) to avoid visual field restrictions. A gradient echo EPI sequence (TR=1 s, TE=55 ms, 36 interleaved slices, resolution=2.3 mm isotropic) with parallel multiband acquisition of 4 simultaneous slices (Breuer *et al.*, 2005) was aligned with the calcarine sulcus and used during hemifield mapping and full field stimulation. An in-plane T1-weighted MPRAGE volume (TR=1.15 ms, TE=3.6 ms, resolution=2×2×2 mm) was acquired to aid registration, as well as a B0 field map (TR=1.17 s, TE_1_=10 ms, TE_2_=14.76 ms) to estimate and correct for local field inhomogeneities. Finally, with the full 32-channel head coil arrangement, a high resolution T1-weighted MPRAGE volume was acquired (TR=2.73 s, TE=3.57 s, resolution=1×1×1 mm).

High-resolution anatomical images were processed with FreeSurfer (v5.3.0, http://www.freesurfer.net) (Dale, Fischl, & Sereno, 1999; Fischl, Sereno, & Dale, 1999), creating a cortical surface render for each participant. A manual definition of the occipital lobe surface was created for each hemisphere in order to restrict data analysis to the occipital cortex. Functional data were pre-processed in SPM12 (Wellcome Trust Centre for Neuroimaging, http://www.fil.ion.ucl.ac.uk/spm). All images were intensity bias-corrected, realigned to the first image of the run and unwarped to correct for movement artefacts and field distortions. A temporal bandpass filter was applied to retain signals between 0.02 and 0.2 Hz. Resulting volumes were registered to the in-plane T1-weighted image and subsequently to the high-resolution T1-weighted image acquired with the full head coil arrangement. Finally, the BOLD time series for each participant were projected onto the individual reconstructed surface by sampling voxels located half-way between the pial and white matter surface in the cortical layer.

Individual HRFs were estimated for each individual participant by sampling data under full field stimulation and averaging the observed time series in the occipital region across trials. This signal was then fitted with a double gamma function (Friston, Frith, Turner, & Frackowiak, 1995), resulting in an individual HRF for each participant’s hemisphere.

### 3.5 pRF modeling

Hemifield mapping runs were analyzed with a forward model pRF approach (Alvarez et al., 2015; Dumoulin & Wandell, 2008), implemented in SamSrf (https://osf.io/2rgsm). MRI data and study analysis code are publicly available at https://osf.io/3mf6n. In brief, model predictions were generated from the *a priori* knowledge of stimulus position at each volume acquired, and under the assumption of an isotropic two-dimensional Gaussian pRF. Predictions were convolved with the individual HRF and compared to the observed signal in a two-stage procedure. First, a coarse fit was conducted by sampling predictions generated from an exhaustive grid of combinations of three pRF parameters (X and Y coordinates, and spatial spread, (σ) and correlating them with a smooth version (FWHM=8.3 mm on the spherical surface mesh) of the observed BOLD time courses. The parameters resulting in the highest correlation at each vertex then formed the starting point for a subsequent fit to the unsmooth data, using a constrained non-linear minimization procedure (Lagarias et al., 1998). Best-fitting model predictions therefore provided estimates of retinotopic location (X and Y coordinates) and pRF size (σ) for each vertex, as well as a scaling factor (β). The coefficient of determination (*R*^*2*^) was taken as the model metric for goodness of fit, and only vertices with *R*^*2*^ > 0.2 were considered in further analysis.

Data resulting from stimulation to each hemifield were analyzed independently, incorporating BOLD time series from clockwise and counter-clockwise runs in each hemisphere. While the stimulated eye differed between participants, all subsequent analyses and discussion refers to hemispheres ipsilateral and contralateral to the stimulated eye, and in the interest of clarity all further illustrations are presented as if stimulation was through the left eye.

An additional analysis was conducted to assess the validity of the dual pRF model of retinotopic encoding in human albinism. Data recorded during stimulation to left and right hemifields were collated and fitted with a) single 2D Gaussian pRF model, as described above and b) dual horizontally-mirrored 2D Gaussian pRF model, which incorporates a second receptive zone of the same size (σ) in the equivalent mirror location across the vertical meridian. Both models were fitted independently for each hemisphere.

### 3.6 Visual area delineation

We manually delineated retinotopic maps on the inflated cortical surface based on polar angle and eccentricity representations derived from the pRF model. Maps were drawn in each hemisphere from data obtained under stimulation to the contralateral visual field. Early visual areas V1, V2 and V3 were reliably identified in all participants, while defining boundaries for further extrastriate areas was variable across participants. Therefore, in order to ensure consistent and sufficient sampling, we restricted our analysis to visual areas V1, V2 and V3.

### 3.7 Data sharing

The conditions of our ethical approval do not permit public archiving of anonymized study data. Researchers seeking access to the data should contact the lead author (Dr Ivan Alvarez), or the local ethics committee (Joint Research and Development Office, Great Ormond Street Hospital for Children NHS Foundation Trust). Access will be granted to named individuals who can demonstrate an appropriate research affiliation, and in accordance with ethical procedures governing the reuse of sensitive data.

## 4 Results

### 4.1 Overlapping representations of ipsilateral and contralateral visual fields

We presented participants with a monocular retinotopic mapping stimulus under two conditions; one stimulating the nasal hemiretina and one stimulating the temporal hemiretina (Figure 1). Responses to nasal and temporal hemiretina stimulation were modelled independently, and pRF estimates obtained for each condition. In control participants, we observed clear lateralization of visual field representations, with the nasal hemiretina represented in the contralateral hemisphere and the temporal hemiretina represented in the ipsilateral hemisphere, as expected (Figure 2). In participants with albinism a contralateral representation of the temporal hemiretina was found to overlap the cortical territory of nasal hemiretina representations in early visual areas, particularly prominent in participants A2 and A3. This confirms the abnormal response lateralization typically seen in albinism and agrees with previous electrophysiological and fMRI studies (Apkarian, Reits, Spekreijse, & Van Dorp, 1983; Dorey, Neveu, Burton, Sloper, & Holder, 2003; von dem Hagen, Hoffmann, & Morland, 2008). Abnormal temporal hemiretina responses in the contralateral hemisphere were largely parafoveal, with the more eccentric representations reverting to the normal uncrossed pattern and appearing in the ipsilateral hemisphere. Substantial overlap was observed in the contralateral hemisphere, in agreement with some animal models of albinism (Guillery, 1986; Guillery et al., 1974; Shatz & LeVay, 1979) and previous fMRI studies (Hoffmann *et al.*, 2003; Kaule *et al.*, 2014). Despite this overlap, no structured representation of polar angle reversals was detected in the abnormal representation (Figure 3).

**Figure 2.**
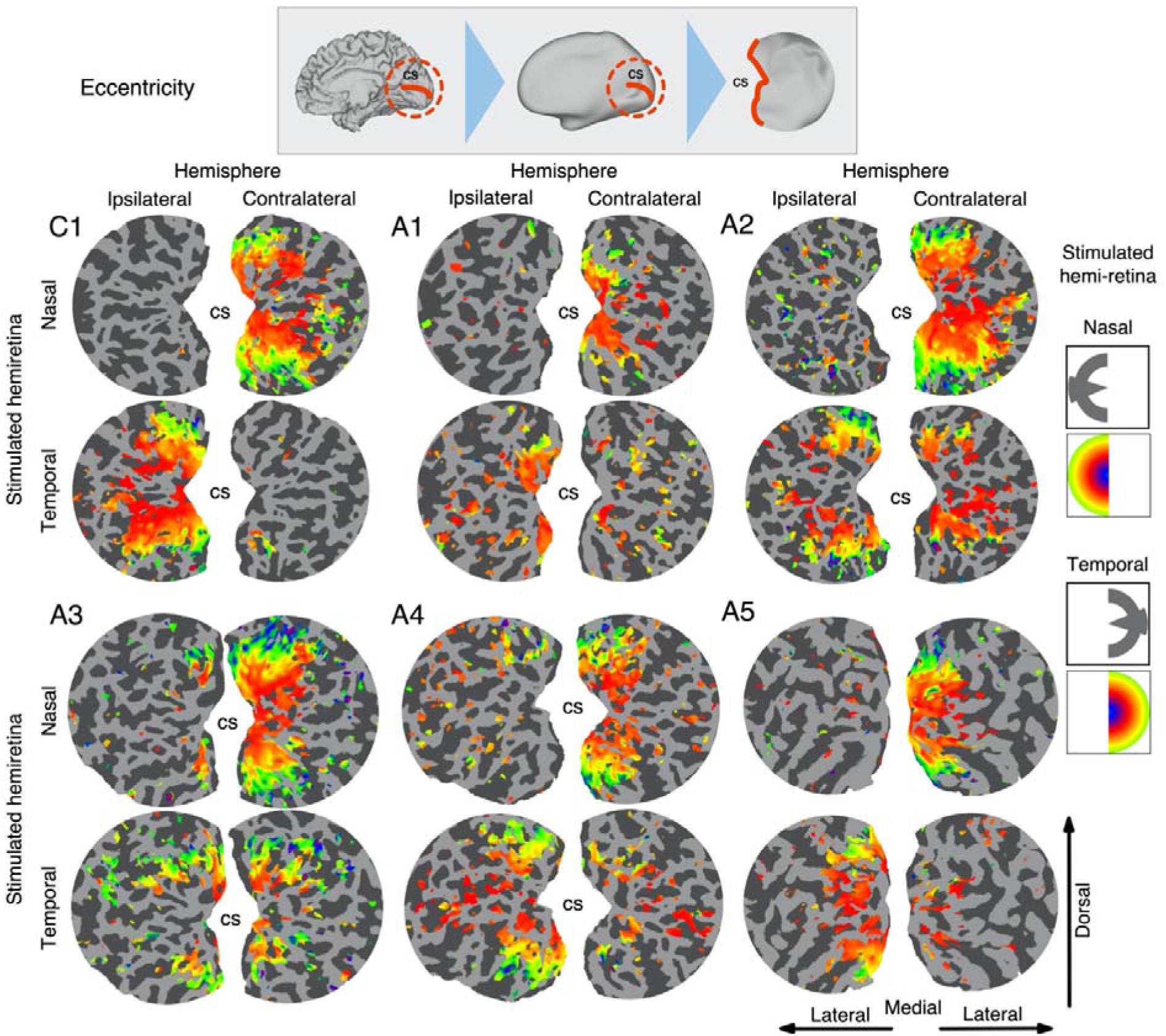
Population receptive field eccentricity on the flattened cortical surface. The cortical surface was inflated, cut and flattened around the calcarine sulcus (cs) for each participant. Stimulation was delivered monocularly to either the nasal or temporal hemiretina, and resulting eccentricity maps are displayed on hemispheres ipsilateral and contralateral to the stimulated eye in five participants with albinism (A1-A5) and one control (C1). In C1, the nasal hemiretina is represented in the contralateral hemisphere, and the temporal hemiretina is represented in the ipsilateral hemisphere. However, visual cortex in A1-A5 showed contralateral responses to temporal hemiretina stimulation. Note the overlap of nasal and temporal hemiretina representations in the contralateral hemisphere. Variability in the vertical line of decussation is apparent from the peripheral extent of the temporal hemiretina representation. Participants received stimulation monocularly to the eye with best visual acuity. Vertices thresholded at R^2^>0.2. cs = calcarine sulcus.

**Figure 3.**
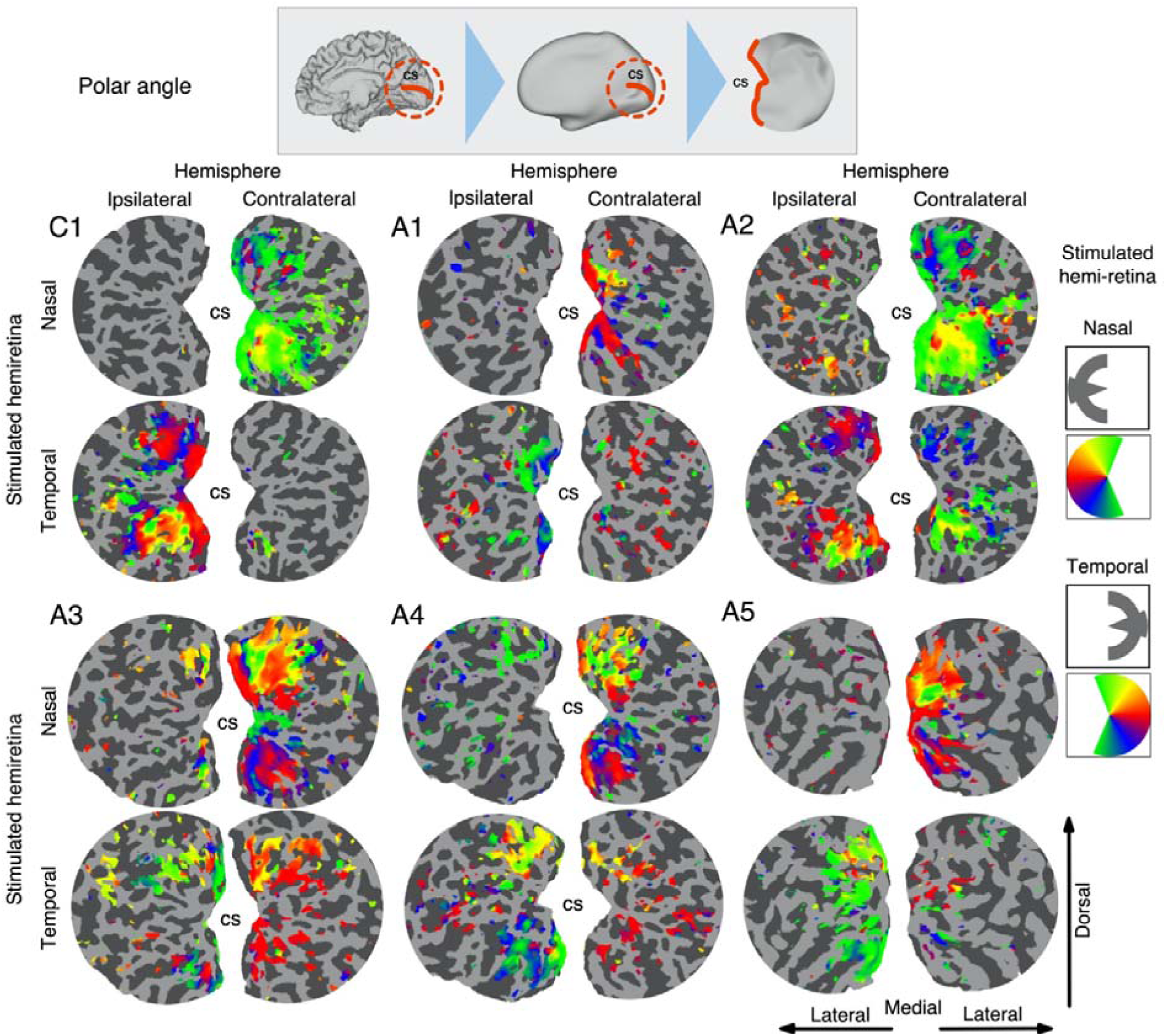
Population receptive field polar angle on the flattened cortical surface. Polar angle maps are displayed on hemispheres ipsilateral and contralateral to the stimulated eye in five participants with albinism (A1-A5) and one control (C1). Polar angle reversals in both dorsal and ventral visual cortex are clear under both nasal stimulation of the contralateral hemisphere and temporal stimulation of the ipsilateral hemisphere. For the abnormal representation of the temporal retina in the contralateral hemisphere in A1-A5, no clear polar angle organization was detected. Vertices thresholded at R^2^>0.2. cs = calcarine sulcus.

An analysis of cortical surface vertices in control participants showed the proportion of early visual cortex (V1, V2 and V3) in the contralateral hemisphere responding to both nasal and temporal hemiretina stimulation was 5.11% (±2.70% SD). The same metric revealed two participants with albinism (A2 = 26.07%, A3 = 18.50%) with significant nasal and temporal response overlaps. No significant spatial overlap was detected for the other participants with albinism (A1 = 6.62%, A4 = 6.11%, A5 = 3.86%).

### 4.2 Nasal bias in receptive field position

Is the representation of the temporal hemiretina abnormally organized in albinotic visual cortex? Answering this question is critical, in order to properly assess the functional role of the abnormal retinal projection. If, for a given cortical point, no systematic discrepancy is found between receptive field positions for the nasal and temporal hemiretina representations, this finding would be consistent with model 1 of albinotic cortical organization, i.e. a contiguous representation of the visual field. On the other hand, if a systematic horizontal bias is detected, with overlapping representations of horizontally distant visual field positions for the nasal and temporal hemiretina on the same cortical territory, it would be consistent with model 2 of albinotic cortical organization.

To answer this question, we examined the estimated pRF locations for the same cortical territory of representational overlap in areas V1, V2 and V3 under two conditions; nasal and temporal hemiretina stimulation. This analysis only included cortical points with significant model fits under both conditions. We then calculated the mean difference in horizontal pRF position between conditions, across visual areas, for each albinotic participant (see Figure 4). This approach revealed a bias for nasal pRFs in the contralateral representation of the temporal hemiretina for all participants with albinism (*mean horizontal shift=*7.90°, *SEM=*4.19°). No compelling evidence emerged for bias towards temporal pRFs in the temporal hemiretina representation. These findings are consistent with model 2 of albinotic cortical organization.

**Figure 4.**
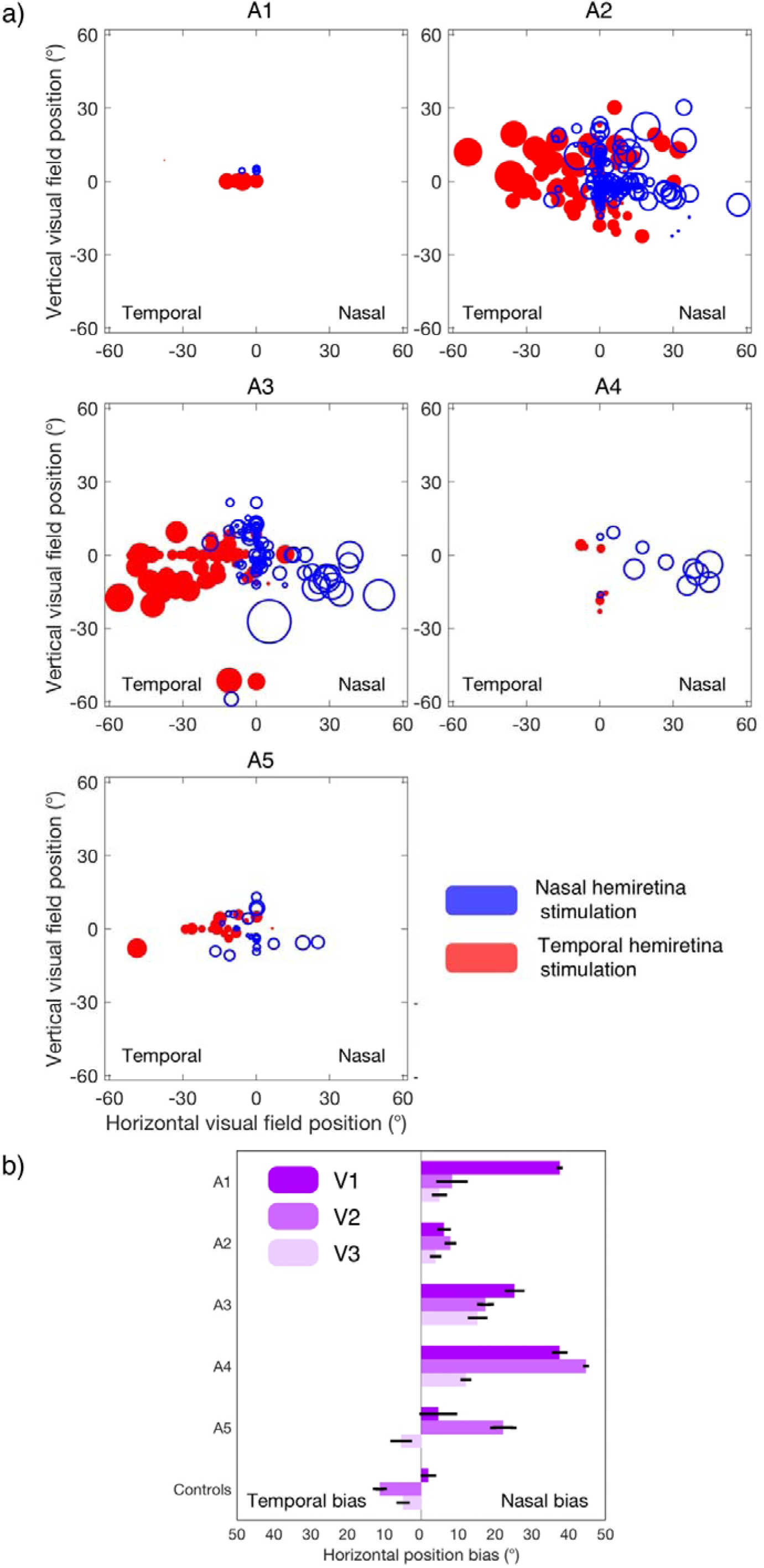
pRF position bias for the same cortical territory under nasal and temporal hemiretina stimulation (a) Back-projection of pRFs in contralateral visual areas V1, V2 and V3 for participants with albinism (A1-A5). Blue circles show pRF positions estimated under nasal hemiretina stimulation, and red circles show pRF positions estimated under temporal hemiretina stimulation. Non-overlap zones indicate a nasal bias in the location of receptive fields in the abnormal temporal hemiretina representation. (b) Horizontal bias in pRF location, as estimated by the mean horizontal position difference between pRF estimates under nasal and temporal hemiretina stimulation, at matching cortical points. Bars extending to the left indicate a bias towards nasal pRFs, and bars extending to the right indicate a bias towards temporal pRFs under temporal hemiretina stimulation, relative to nasal hemiretina stimulation. All participants with albinism show bias towards a nasal shift in the abnormal temporal hemiretina representation in the hemisphere contralateral to stimulation. Error bars indicate SEM.

### 4.3 Variability in receptive field size

In addition to biases in pRF location, another potential indicator for altered functional roles in cortical processing may be enlarged or constricted pRF sizes in the abnormal retinal representation zone. To examine this, the significant vertices for all subjects in each group were collated and fitted with a linear model (Figure 5). Following stimulation to the temporal hemiretina, both controls and participants with albinism displayed a positive monotonic relationship between pRF size and eccentricity in the hemisphere ipsilateral to stimulation. In order to assess these trends statistically, individual participant estimates of pRF size were summarized by averaging values into 30 bins, each spanning 2° of eccentricity. Binned pRF sizes were introduced to a mixed ANOVA model as the dependent variable, with group (control, albinotic) as the between-subject independent variable, and both pRF eccentricity and visual area (V1, V2, V3) as within-subject independent variables. No statistically significant interaction between group and eccentricity was detected (*F*(1, 13) = 0.01, *p* = 0.948, η^2^ = 10^−3^), nor significant 3-way interaction between group, visual area and eccentricity (*F*(2, 26) = 0.84, *p* = 0.445, η^2^ = 0.06). The individual-level trends in pRF size were assessed by comparing estimates of binned pRF size in single albinotic participants against the control group mean estimate in visual areas V1, V2 and V3 (see Figure 6). The contralateral response to nasal retina stimulation and the ipsilateral response to temporal retina stimulation were found to be broadly similar between albinotic participants and controls, albeit with high inter-individual variability in the albinotic group. These results indicate the abnormal temporal hemiretina representation in the contralateral hemisphere follows the same organization principles at the group level, as regularly organized cortex in both controls and participants with albinism.

**Figure 5.**
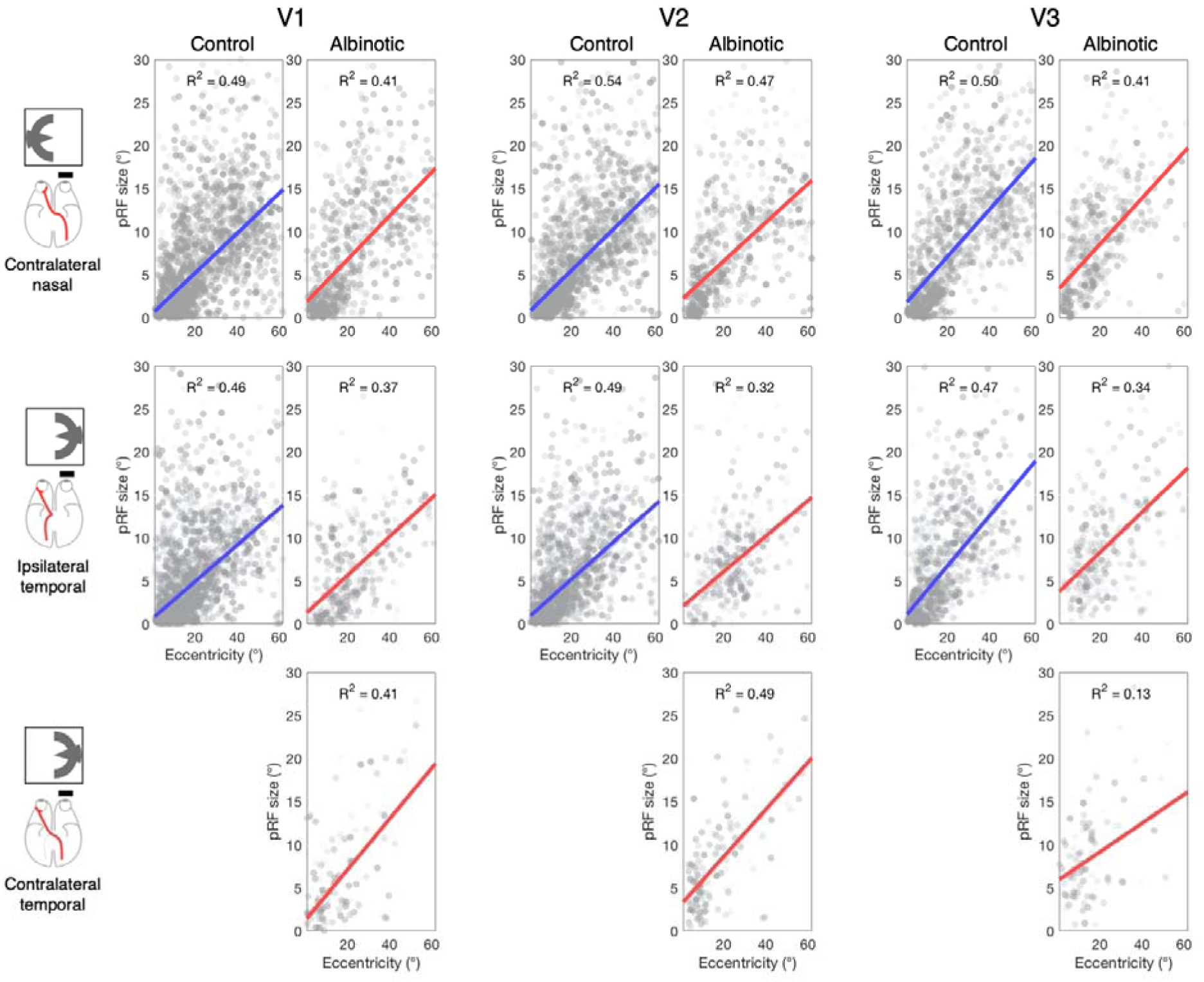
pRF size as a function of receptive field eccentricity in cortical visual areas V1, V2 and V3 for controls (N=10, blue) and albinotic (N=5) participants. Best-fitting linear model (blue = control, red = albinotic) and matching coefficient of determination (R^2^) shown for each condition. Both groups show a monotonic increase in pRF size with eccentricity. Contralateral responses to temporal retina stimulation omitted for controls, as no significant responses were detected under that condition. No significant differences between groups in the relationship between pRF size and eccentricity were observed.

**Figure 6.**
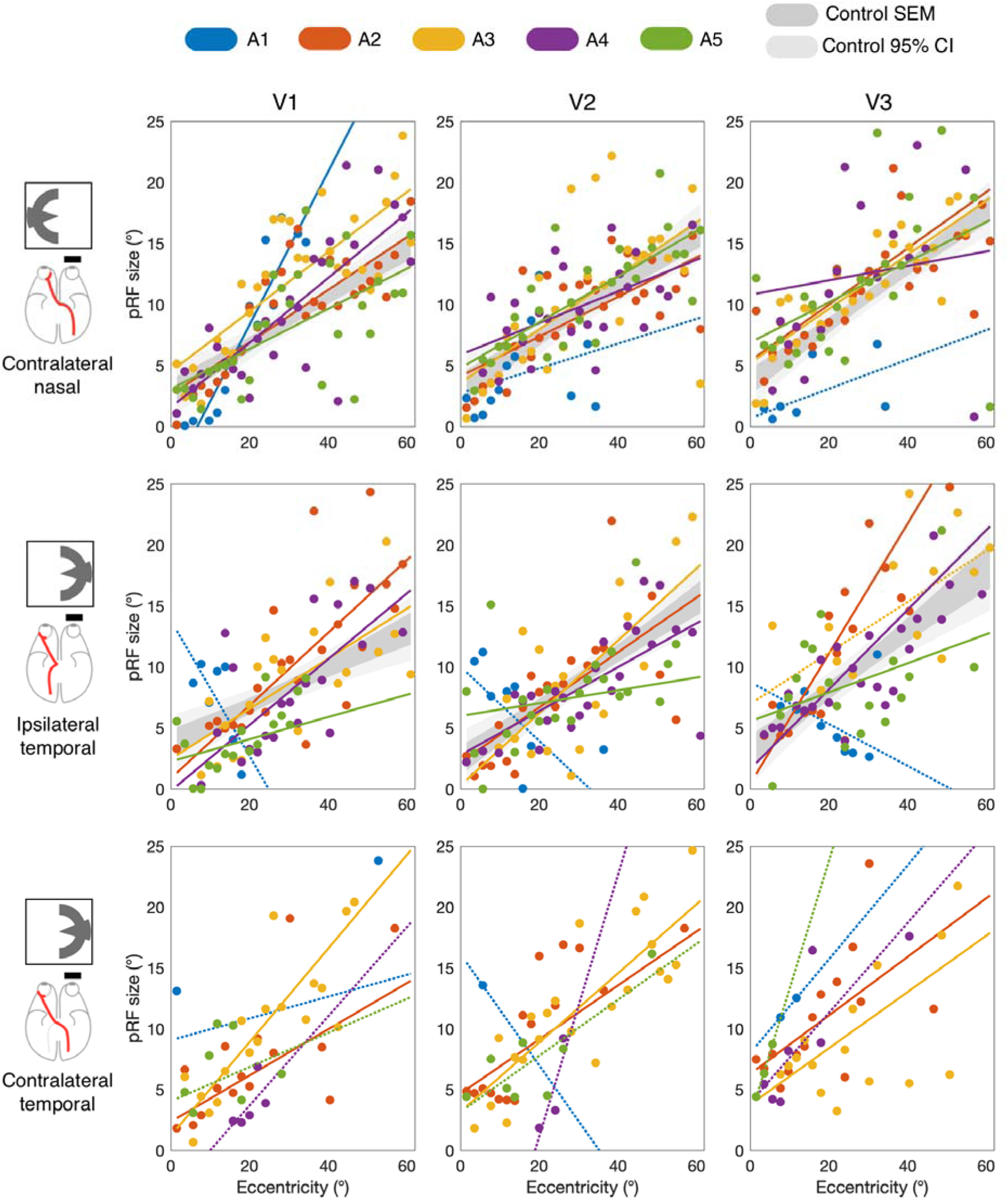
pRF size as a function of receptive field eccentricity in cortical visual areas V1, V2 and V3 for individual participants. Mean pRF size in successive 2° eccentricity bins and best linear fits displayed for participants with albinism (A1-A5). Control data (N=10) shown as standard error of the mean (dark grey) and 95% confidence intervals (light grey) of binned pRF sizes. A1-A5 display trends consistent with control data in pRF size increase with eccentricity in the contralateral representation of both nasal and temporal retina. Dashed lines indicate insufficient sampling of significant vertices across eccentricity (<50% bins) to fit full regression model. No normative data displayed in bottom row as no significant contralateral responses to temporal retina stimulation were detected in controls.

### 4.4 No evidence for dual receptive fields

In order to further explore the underlying organization of visual receptive fields in albinotic visual cortex, an additional analysis was performed where responses to nasal and temporal hemiretina stimulation were modeled using both a classic 2D Gaussian pRF model, as implemented in the previous section, and alternatively with a dual pRF model, with two receptive fields, mirror-symmetric across the vertical median, as a neuronal population-level approximation of the dual receptive field model (Klemen *et al.*, 2012). The single and dual pRF models were fitted independently across early visual cortex. To assess differences in fits between the two models, we compared goodness of fit (R^2^) estimates for each participant with a series of matched samples tests. As the distribution of R^2^ were non-normal (Figure 7), we used the non-parametric Wilcoxon signed-rank test. Bonferroni correction for multiple comparison was applied across all tests conducted. For all participants, we observed similar or significantly larger R^2^ estimates under the single pRF model when compared to the dual pRF model. Significantly better fits for the single pRF model were detected for the control group in all visual areas; V1 *(Z* = 36.88, *p*_*corr*_ = 10^−296^, effect size *r* = 0.62), V2 (Z = 26.31, *p*_*corr*_ = 10^−151^, effect size *r* = 0.52) and V3 (*Z* = 17.59, *p*_*corr*_ = 10^−68^, effect size *r* = 0.40). For participants with albinism, significantly better single pRF model fits were detected in V1 for A1, A2, A3 and A4, in V2 for A2, A3, A4, and in V3 for A2, A3 and A4 (all tests *p*_corr_ < 0.01). Overall, individual results show the dual receptive field model is either statistically indistinguishable, or a significantly poorer fit than the single receptive field throughout the cortical areas tested. While the overall pattern of responses across A1-A5 favored the classic single receptive field model, a further line of enquiry was pursued to assess whether the dual receptive field was a better fit in spatially localized subsets of vertices within the visual areas sampled. The proportion of vertices where the dual receptive field model provided a better fit (*R*^*2*^ *difference* > 0.1) was consistently below 9% across all participants in bilateral V1, V2 and V3 vertices (A1=0.22%, A2=9.08%, A3=8.27%, A4=2.18%, A5=2.34%). In the small proportion of vertices where the dual pRF model outperformed the single pRF model, the mean difference in goodness of fit was small (*mean R*^*2*^ *difference*; A1=0.05, A2=0.13, A3=0.13, A4=0.06, A5=0.07). In addition, the spatial distribution of this subset of vertices revealed no coherent structure in visual cortical areas (Figure 8). Together, these results indicate that the dual receptive field is a poor model of the overlapping cortical representation in contralateral visual cortex in participants with albinism.

**Figure 7.**
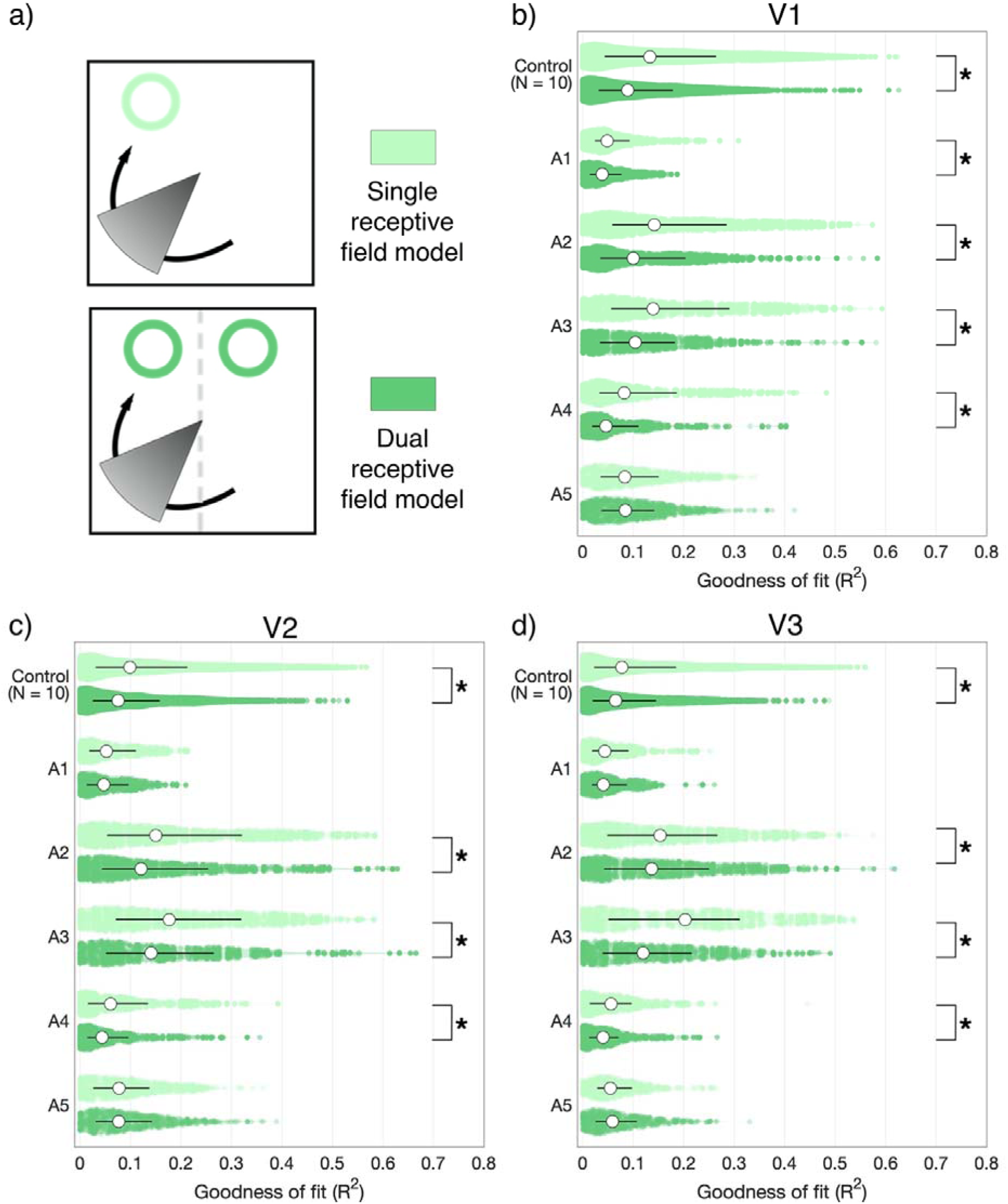
Single vs. dual pRF model fits. (a) BOLD responses to bilateral visual field stimulation were fitted with either a single or dual pRF model. Dual receptive fields were mirrored across the vertical meridian, and of equal spread (σ). (b-d) Distribution of goodness of fit (R^2^) estimates for each model are displayed in areas V1, V2 and V3 for five participants with albinism (A1-A5) and 10 healthy controls. The single pRF model either outperformed the dual pRF model or was statistically equivalent for all participants in V1, V2 and V3. No individual comparison yielded significantly better fits for the dual pRF model over the single pRF model. Asterisks denote significant Wilcoxon signed-rank tests *p*<0.01, corrected for multiple comparisons.

**Figure 8.**
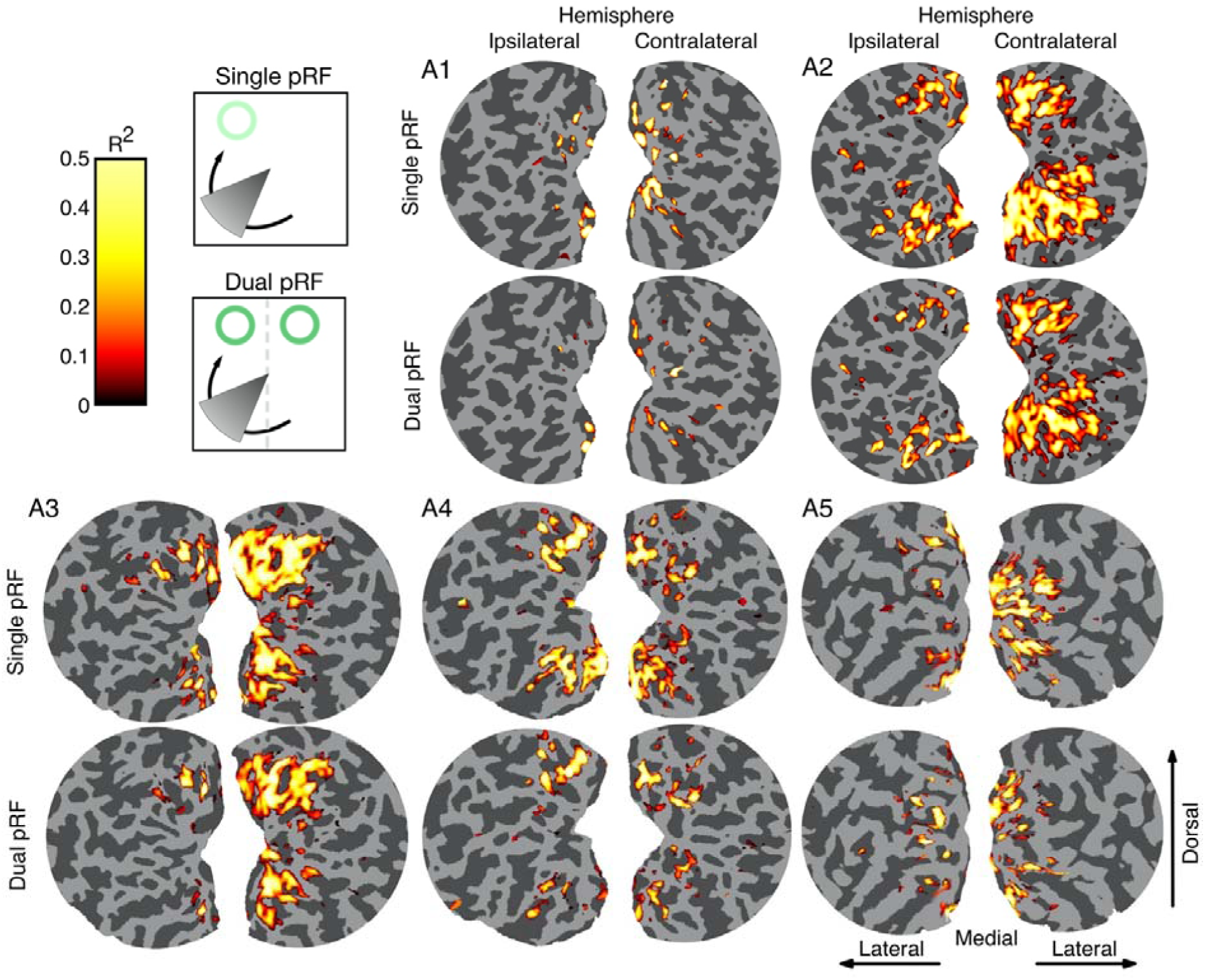
Spatial distribution of significant vertices in single and dual pRF models. Model goodness of fit (R^2^) is shown on the flattened occipital cortex for five participants with albinism (A1-A5), on hemispheres ipsilateral (left column) and contralateral (right column) to monocular stimulation. For each participant, the top row shows cortical locations where the single pRF model resulted in significant models fits, and the bottom row where the dual pRF models resulted in significant model fits (R^2^ > 0.2). Vertices where the dual pRF model outperforms the single pRF model do not form spatially coherent clusters and represent <9% of significant vertices across visual areas V1, V2 and V3.

## 5 Discussion

We identified and characterized the abnormal visual field representation in five rare participants with albinism and no nystagmus. Following monocular stimulation of the temporal hemiretina, significant BOLD responses were observed in both ipsilateral and contralateral occipital cortices, reproducing the widely reported asymmetric lateralization of responses in albinism (Apkarian *et al.*, 1983; Carroll *et al.*, 1980; Creel *et al.*, 1974; Hagen *et al.*, 2008; Hedera *et al.*, 1994). The abnormal response was found to be retinotopically organized, with the temporal hemiretina representation largely overlapping the nasal representation in the contralateral hemisphere, in agreement with previous electrophysiological (Guillery *et al.*, 1984) and fMRI findings (Hagen, Houston, Hoffmann, Jeffery, & Morland, 2005; Hoffmann et al., 2003; Kaule et al., 2014; Morland et al., 2002). This abnormal temporal representation was characterized by a shifted line of decussation, with foveal portions represented contralaterally, and eccentric portions represented in the ipsilateral hemisphere. This is again consistent with previous human fMRI literature (Hagen, Houston, Hoffmann, & Morland, 2007; Hoffmann et al., 2003; Kaule et al., 2014).

One advantage of performing visual mapping with participants with unimpaired fixation is the potential for careful mapping of receptive field properties. In this study, we implemented a hemifield-restricted stimulus where in each run, a pair of de-phased, contrast-reversing apertures were presented to a single hemifield while participants fixated, therefore exclusively stimulating either the nasal or temporal hemiretina.

Three findings of interest emerge from the analysis presented here. First, no evidence for systematically altered pRF sizes point towards a conservative role for visual areas representing the abnormally overlapping nasal and temporal hemiretina representations. The lack of altered pRF properties in area V1 is surprising given previous reports of reduced calcarine fissure length (Neveu & Jeffery, 2007) and increased grey matter volume and cortical thickness in albinotic V1 (Bridge *et al.*, 2014; Hagen *et al.*, 2005). While one may expect such structural abnormalities to be linked with concomitant functional differences such as receptive field spatial sensitivity, as approximated by pRF metrics, it is worth highlighting that albinism is a heterogeneous disorder (Carroll et al., 1980; Neveu, Jeffery, Burton, Sloper, & Holder, 2003). It is therefore possible that individuals with no nystagmus in the primary position, such as those reported on here, may represent a unique population with reduced chiasmal abnormalities and therefore reduced anatomical or functional consequences at the level of cortical representations.

Non-altered receptive field properties in albinism raise the question of functional significance for overlapping visual field representations in contralateral cortex. While it was originally postulated that the abnormal temporal representation might be suppressed (Guillery *et al.*, 1974; Kaas, 2005), electrophysiological and functional imaging evidence in humans suggests this abnormal representation is indeed encoding valuable retinotopic information (Kaule *et al.*, 2014). Here we show that both overlapping responses in contralateral V1 display similar pRF size increases with eccentricity, indicating a similar functional role in encoding of retinotopic information at the earliest stage of cortical processing. Significant levels of activation, retinotopic organization and coherent receptive field increases with eccentricity all indicate this region plays an active role in the representation of the temporal hemiretina. It is also worth noting that inspection of responses to temporal hemiretina stimulation reveals a split representation across the cortical hemispheres, rather than a duplicate representation. This highlights the importance of the abnormal temporal representation in maintaining continuity of the spatial representation of the visual field, which may not be afforded by a suppressed representation.

How is visual information encoded when overlapping retinotopic representations are present in albinism? One possibility is that cortical cells with conservative organization represent all the retinal inputs they receive, giving rise to dual receptive field cells. Such cells would be responsive to mirrored nasal and temporal retinal locations and non-discriminatory of originating hemifield. Imaging evidence has suggested such cells may be present in achiasma, a related condition of optic nerve misrouting (Hoffmann *et al.*, 2012). Alternatively, hemifield discrimination can take place at the early cortical level, with single receptive field cells responding to a single retinal location in order to integrate ocular dominance column information. Here, we modeled BOLD responses to bilateral visual field stimulation in participants with albinism with two models; a single receptive field model with a single pRF zone and a dual receptive field model with two mirror-symmetrical pRFs divided by the vertical meridian. While other models of multiple spatial encoding are possible, this model was chosen in order to assess the explicit hypothesis of horizontally-mirrored receptive fields postulated by (Klemen *et al.*, 2012). When assessed for model evidence, the single receptive field model outperformed the dual receptive field model in all participants, both control and participants with albinism. Vertices where the dual receptive field model provided a better fit were fewer than 9% of those sampled, and displayed no spatial clustering in cortex. Overall, evidence in this study suggests a rejection of the dual receptive field encoding model for human albinism, in agreement with previous psychophysical evidence (Klemen *et al.*, 2012). These results, therefore, suggest hemifield information is segregated at an early cortical level. Given the normal or sub-normal stereovision reported in some patients with albinism (Lee et al., 2001; Summers, 1996), successful integration of ocular dominance column information can occur without major sensory conflict (cf. (Tsytsarev, Arakawa, Zhao, Chédotal, & Erzurumlu, 2017). In order to achieve this, a degree of cortical plasticity must come into play, as contralateral cortical territories normally represent retinotopically mirror-equivalent positions in left and right visual fields that must be functionally segregated. If the ‘true albino’ pattern (Guillery *et al.*, 1984) is an accurate model of visual field representation in the human, and the present study as well as recent fMRI findings suggest that may be the case (Hagen et al., 2007; Hoffmann et al., 2003; Kaule et al., 2014; Wolynski, Kanowski, Meltendorf, Behrens-Baumann, & Hoffmann, 2010), hemifield segregation may be achieved through the presence of hemifield columns. Nevertheless, the correct selection of sensory information across hemifield columns and between ocular dominance columns for stereopsis is not easily achieved with a conservative retinotopic organization. Instead, these findings suggest a degree of developmental plasticity involved in providing a disambiguated sensory input to early visual processing, leading to normal or near-normal visual perception in human albinism.

## Abbreviations

BOLD: blood-oxygen-level dependent;
fMRI: functional magnetic resonance imaging;
HRF: hemodynamic response function;
pRF: population receptive field

## 6 Appendices

### Appendix A.

Monocular visual field perimetry for participants with albinism (A1-A5). Perimetry for outer targets (I4e) are displayed in blue, overlaid on the mean perimeter (black) and standard deviation (grey) of control participants (N=10). Target speed 5°/s. Outer perimeter for A1-A5 fall largely within the normal control perimeter, excepting a reduction in the nasal visual field in A2 and dorsal visual field for A4. During fMRI, monocular stimulation was delivered to the right eye for A2 and left for A1, A3, A4 and A5.

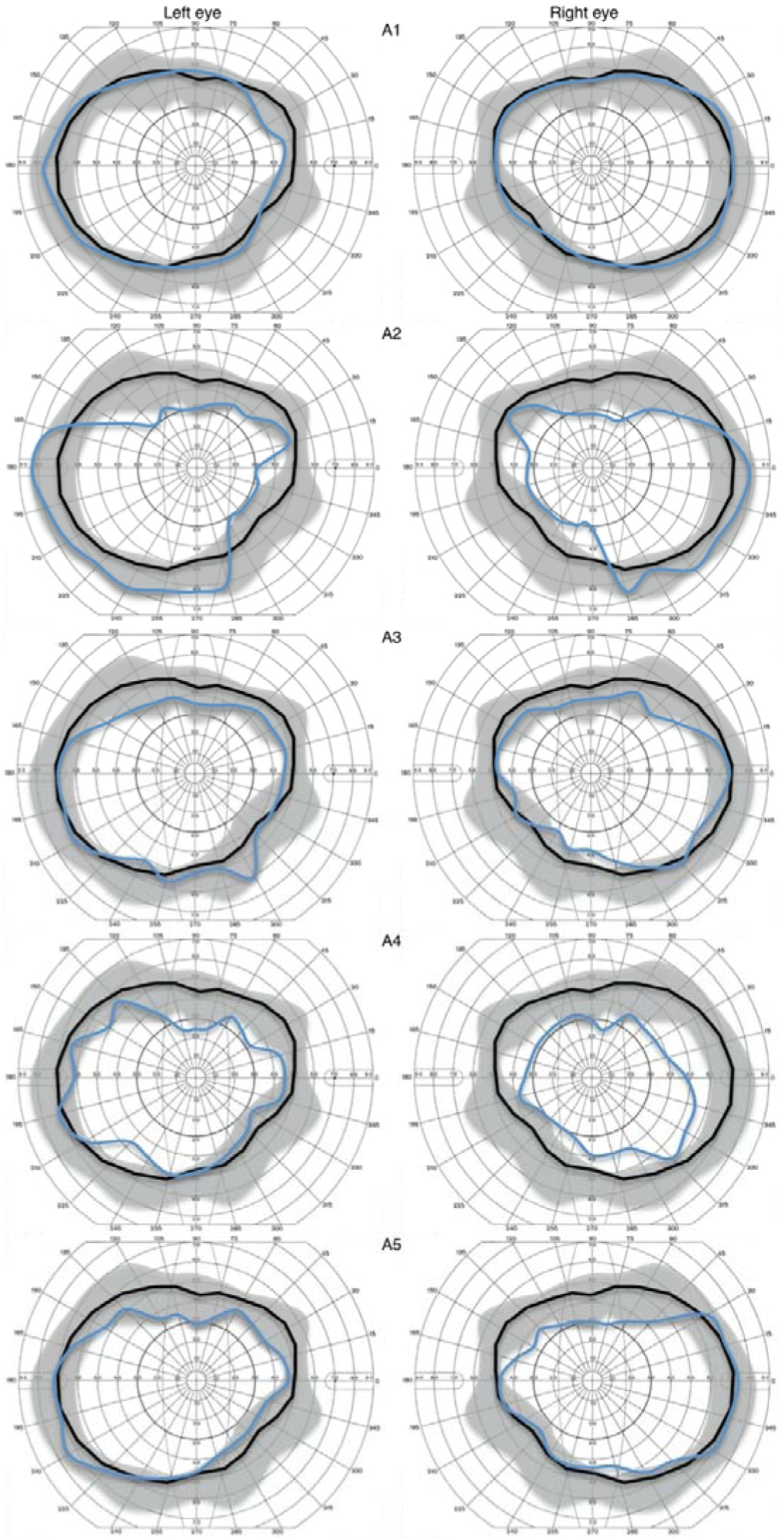

### Appendix B.

Optical coherence tomography (OCT) of the retina in participants with albinism (A1-A5). Transverse location and three axial slices are shown for the left and right eyes. Retinal scans were centred on the macula, with no clear foveal pit displayed (foveal hypoplasia), a characteristic of albinism

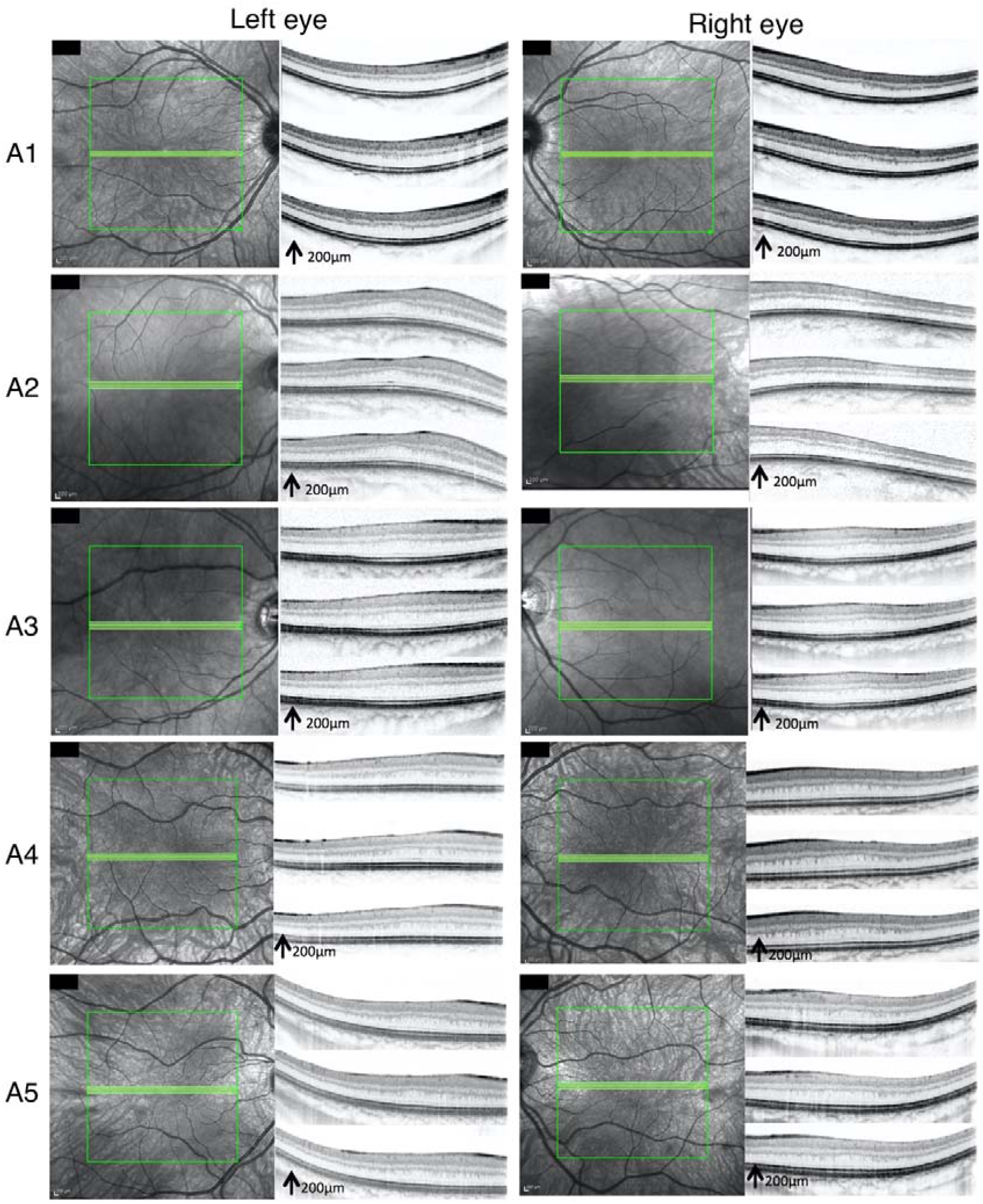

## 7 Acknowledgements

We thank Prof Anthony Moore, Prof Glen Jeffery, Dr Magella Neveu and Dr Richard Bowman, and the ophthalmology teams at Moorfields Eye Hospital and Great Ormond Street Hospital for their assistance during participant recruitment. We also thank Dr Benjamin de Haas and Dr Christina Moutsiana for their assistance during data acquisition and valuable discussions. No part of the study procedures or analyses was pre-registered prior to the research being conducted.

## 8 Funding

This work was supported by a UCL SLMS Grand Challenge Studentship in Biomedicine, funded by the UCL School for Life and Medical Sciences, UCLH/UCL Comprehensive Biomedical Research Centre and the Specialist Biomedical Research Centres at Moorfields/UCL and Great Ormond Street/UCL.

## Notes

### Competing Interest Statement

The authors have declared no competing interest.

### Summary of Updates

Manuscript accepted for publication.

## References

Alvarez, I., de Haas, B., Clark, C. A., Rees, G., & Schwarzkopf, D. S. (2015). Comparing different stimulus configurations for population receptive field mapping in human fMRI. Frontiers in Human Neuroscience, 9, 96. http://doi.org/10.3389/fnhum.2015.00096/abstract

Apkarian, P., & Shallo-Hoffmann, J. (1991). VEP projections in congenital nystagmus; VEP asymmetry in albinism: A comparison study. Investigative Ophthalmology & Visual Science, 32(9), 2653–2661.

Apkarian, P., Reits, D., Spekreijse, H., & Van Dorp, D. (1983). A decisive electrophysiological test for human albinism. Electroencephalography and Clinical Neurophysiology, 55(5), 513–531. http://doi.org/10.1016/0013-4694(83)90162-1

Binda, P., Thomas, J. M., Boynton, G. M., & Fine, I. (2013). Minimizing biases in estimating the reorganization of human visual areas with BOLD retinotopic mapping. Journal of Vision, 13(7), 13–13. http://doi.org/10.1167/13.7.13

Brainard, D. H. (1997). The psychophysics toolbox. Spatial Vision, 10(4), 433–436.

Bressler, D. W., & Silver, M. A. (2010). Spatial attention improves reliability of fMRI retinotopic mapping signals in occipital and parietal cortex. NeuroImage, 53(2), 526–533. http://doi.org/10.1016/j.neuroimage.2010.06.063

Breuer, F. A., Blaimer, M., Heidemann, R. M., Mueller, M. F., Griswold, M. A., & Jakob, P. M. (2005). Controlled aliasing in parallel imaging results in higher acceleration (CAIPIRINHA) for multi-slice imaging. Magnetic Resonance in Medicine, 53(3), 684–691. http://doi.org/10.1002/mrm.20401

Bridge, H., Hagen, von dem E. A. H., Davies, G., Chambers, C., Gouws, A., Hoffmann, M., & Morland, A. B. (2014). Changes in brain morphology in albinism reflect reduced visual acuity. Cortex, 56, 64–72. http://doi.org/10.1016/j.cortex.2012.08.010

Carroll, W. M., Jay, B. S., McDonald, W. I., & Halliday, A. M. (1980). Two distinct patterns of visual evoked response asymmetry in human albinism. Nature, 286(5773), 604–606. http://doi.org/10.1038/286604a0

Charles, S. J., Green, J. S., Grant, J. W., Yates, J., & Moore, A. T. (1993). Clinical features of affected males with X-linked ocular albinism. British Journal of Ophthalmology, 77(4), 222–227.

Cooper, M. L., & Blasdel, G. G. (1980). Regional variation in the representation of the visual field in the visual cortex of the Siamese cat. Journal of Comparative Neurology, 193(1), 237–253. http://doi.org/10.1002/cne.901930116

Creel, D., Witkop, C. J., & King, R. A. (1974). Asymmetric visually evoked potentials in human albinos: Evidence for visual system anomalies. Investigative Ophthalmology, 13(6), 430–440.

Crossland, M. D., Morland, A. B., Feely, M. P., Hagen, von dem E. A. H., & Rubin, G. S. (2008). The effect of age and fixation instability on retinotopic mapping of primary visual cortex. Investigative Ophthalmology & Visual Science, 49(8), 3734–3739. http://doi.org/10.1167/iovs.07-1621

Dale, A. M. A., Fischl, B. B., & Sereno, M. I. M. (1999). Cortical surface-based analysis - I. segmentation and surface reconstruction. NeuroImage, 9(2), 179–194. http://doi.org/10.1006/nimg.1998.0395

Dorey, S. E., Neveu, M. M., Burton, L. C., Sloper, J. J., & holder, G. E. (2003). The clinical features of albinism and their correlation with visual evoked potentials. British Journal of Ophthalmology, 87(6), 767–772.

Dumoulin, S. O., & Wandell, B. A. (2008). Population receptive field estimates in human visual cortex. NeuroImage, 39(2), 647–660. http://doi.org/10.1016/j.neuroimage.2007.09.034

Fischl, B., Sereno, M. I., & Dale, A. M. (1999). Cortical surface-based analysis. II: Inflation, flattening, and a surface-based coordinate system. NeuroImage, 9(2), 195–207. http://doi.org/10.1006/nimg.1998.0396

Friston, K. J., Frith, C. D., Turner, R., & Frackowiak, R. S. (1995). Characterizing evoked hemodynamics with fMRI. NeuroImage, 2(2), 157–165. http://doi.org/10.1006/nimg.1995.1018

Guillery, R. W. (1986). Neural abnormalities of albinos. Trends in Neurosciences, 9(8), 364–367. http://doi.org/10.1016/0166-2236(86)90115-3

Guillery, R. W., Casagrande, V. A., & Oberdorfer, M. D. (1974). Congenitally abnormal vision in Siamese cats. Nature, 252(5480), 195–199. http://doi.org/10.1038/252195a0

Guillery, R. W., Hickey, T. L., Kaas, J. H., Felleman, D. J., Debruyn, E. J., & Sparks, D. L. (1984). Abnormal central visual pathways in the brain of an albino green monkey (cercopithecus aethiops). Journal of Comparative Neurology, 226(2), 165–183. http://doi.org/10.1002/cne.902260203

Guillery, R. W., Okoro, A. N., & Witkop, C. J. (1975). Abnormal visual pathways in the brain of a human albino. Brain Research, 96(2), 373–377. http://doi.org/10.1016/0006-8993(75)90750-7

Hagen, von dem E. A. H., Hoffmann, M. B., & Morland, A. B. (2008). Identifying human albinism: A comparison of VEP and fMRI. Investigative Ophthalmology & Visual Science, 49(1), 238–249. http://doi.org/10.1167/iovs.07-0458

Hagen, von dem E. A. H., Houston, G. C. G., Hoffmann, M. B. M., Jeffery, G. G., & Morland, A. B. (2005). Retinal abnormalities in human albinism translate into a reduction of grey matter in the occipital cortex. European Journal of Neuroscience, 22(10), 2475–2480. http://doi.org/10.1111/j.1460-9568.2005.04433.x

Hagen, von dem E. A. H., Houston, G. C., Hoffmann, M. B., & Morland, A. B. (2007). Pigmentation predicts the shift in the line of decussation in humans with albinism. European Journal of Neuroscience, 25(2), 503–511. http://doi.org/10.1111/j.1460-9568.2007.05303.x

Hedera, P., Lai, S., Haacke, E. M., Lerner, A. J., Hopkins, A. L., Lewin, J. S., & Friedland, R. P. (1994). Abnormal connectivity of the visual pathways in human albinos demonstrated by susceptibility-sensitized MRI. Neurology, 44(10), 1921–1926.

Hoffmann, M. B., & Dumoulin, S. O. (2015). Congenital visual pathway abnormalities: a window onto cortical stability and plasticity. Trends in Neurosciences, 38(1), 55–65. http://doi.org/10.1016/j.tins.2014.09.005

Hoffmann, M. B., Kaule, F. R., Levin, N., Masuda, Y., Kumar, A., Gottlob, I., et al. (2012). Plasticity and stability of the visual system in human achiasma. Neuron, 75(3), 393–401. http://doi.org/10.1016/j.neuron.2012.05.026

Hoffmann, M. B., Tolhurst, D. J., Moore, A. T., & Morland, A. B. (2003). Organization of the visual cortex in human albinism. Journal of Neuroscience, 23(26), 8921–8930.

Hubel, D. H., & Wiesel, T. N. (1971). Aberrant visual projections in the Siamese cat. Journal of Physiology, 218(1), 33–62.

Hummer, A., Ritter, M., Tik, M., Ledolter, A., Woletz, M., holder, G. E., et al. (2016). Eyetracker-based gaze correction for robust mapping of population receptive fields. NeuroImage, 1–38. http://doi.org/10.1016/j.neuroimage.2016.07.003

Kaas, J. H. (2005). Serendipity and the Siamese cat: The discovery that genes for coat and eye pigment affect the brain. ILAR Journal, 46(4), 357–363.

Kaas, J. H., & Guillery, R. W. (1973). The transfer of abnormal visual field representations from the dorsal lateral geniculate nucleus to the visual cortex in siamese cats. Brain Research, 59, 61–95. http://doi.org/10.1016/0006-8993(73)90253-9

Kaule, F. R., Wolynski, B., Gottlob, I., Stadler, J., Speck, O., Kanowski, M., et al. (2014). Impact of chiasma opticum malformations on the organization of the human ventral visual cortex. Human Brain Mapping, 35(10), ka–5105. http://doi.org/10.1002/hbm.22534

Kinnear, P. E. P., Jay, B. B., & Witkop, C. J. C. (1985). Albinism. Survey of Ophthalmology, 30(2), 75–101.

Klemen, J., Hoffmann, M. B., & Chambers, C. D. (2012). Cortical plasticity in the face of congenitally altered input into V1. Cortex, 48(10), 1362–1365. http://doi.org/10.1016/j.cortex.2012.03.012

Lagarias, J. C., Reeds, J. A., Wright, M. H., & Wright, P. E. (1998). Convergence properties of the Nelder-Mead simplex method in low dimensions. SIAM Journal on Optimization, 9(1), 112–147.

Lee, K. A., King, R. A., & Summers, C. G. (2001). Stereopsis in patients with albinism: Clinical correlates. Journal of American Association for Pediatric Ophthalmology and Strabismus, 5(2), 98–104. http://doi.org/10.1067/mpa.2001.112441

Levin, N., Dumoulin, S. O., Winawer, J., Dougherty, R. F., & Wandell, B. A. (2010). Cortical maps and white matter tracts following long period of visual deprivation and retinal image restoration. Neuron, 65(1), 21–31. http://doi.org/10.1016/j.neuron.2009.12.006

Morland, A. B., Hoffmann, M., Neveu, M. M., & Holder, G. E. (2002). Abnormal visual projection in a human albino studied with functional magnetic resonance imaging and visual evoked potentials. Journal of Neurology, Neurosurgery & Psychiatry, 72(4), 523–526.

Neveu, M. M., & Jeffery, G. G. (2007). Chiasm formation in man is fundamentally different from that in the mouse, 21(10), 1264–1270. http://doi.org/10.1038/sj.eye.6702839

Neveu, M. M., Jeffery, G. G., Burton, L. C., Sloper, J. J., & Holder, G. E. (2003). Age-related changes in the dynamics of human albino visual pathways. European Journal of Neuroscience, 18(7), 1939–1949. http://doi.org/10.1046/j.1460-9568.2003.02929.x

Pelli, D. G. D. (1997). The VideoToolbox software for visual psychophysics: Transforming numbers into movies. Spatial Vision, 10(4), 437–442.

Shatz, C. J., & LeVay, S. (1979). Siamese cat: altered connections of visual cortex. Science, 204(4390), 328–330.

Summers, C. G. (1996). Vision in albinism. Transactions of the American Ophthalmological Society, 94, 1095–1155.

Tsytsarev, V., Arakawa, H., Zhao, S., Chédotal, A., & Erzurumlu, R. S. (2017). Behavioral Consequences of a Bifacial Map in the Mouse Somatosensory Cortex. Journal of Neuroscience, 37(30), 7209–7218. http://doi.org/10.1523/JNEUROSCI.0598-17.2017

Wandell, B. A., & Winawer, J. (2015). Computational neuroimaging and population receptive fields. Trends in Cognitive Sciences, 19(6), 349–357. http://doi.org/10.1016/j.tics.2015.03.009

Wolynski, B., Kanowski, M., Meltendorf, S., Behrens-Baumann, W., & Hoffmann, M. B. (2010). Self-organisation in the human visual system-Visuo-motor processing with congenitally abnormal V1 input. Neuropsychologia, 48(13), 3834–3845. http://doi.org/10.1016/j.neuropsychologia.2010.09.011

